# STAMPS: Signal-peptide Transformer for Augmenting Mammalian Protein Secretion

**DOI:** 10.1101/2025.11.20.689577

**Authors:** Jacopo Gabrielli, Marino Exposito-Rodriguez, Justina Briliūtė, Matthew Burrell, George Thom, Cleo Kontoravdi, Francesca Ceroni

## Abstract

While secretion plays a key role in the diverse applications of cell engineering, to date only a handful of mammalian signal peptides have been characterised in depth and systematic efforts to build novel variants remain sporadic. We present STAMPS (Signal-peptide Transformer for Augmenting Mammalian Protein Secretion), a generative, autoregressive transformer fine-tuned on ∼6,000 mammalian signal peptides to design *de novo* sequences that modulate and enhance secretion across proteins and hosts. We show that STAMPS can be used to identify candidate signal peptides that outperform the widely used IgG κ light-chain leader (IgKL) when used to secrete EGFP in HEK293T and CHO cells, and hEPO in HEK293T cells. When incorporated in an industrial cell line development framework, STAMPS leads to the generation of signal peptides that yield ∼2.3-fold gain in secretion of a VHH-Fc compared to the internal industrial benchmark in CHO G22 cells, with the same candidates ranking highly in both CHO and HEK293T hosts. Sequence-to–function analysis highlights a longer, strongly hydrophobic core and a tightly positioned cleavage site as drivers of this strong performance. Together, these results establish data-driven, generative design of mammalian signal peptides as a practical route to tune and improve secretion for bioproduction and cell engineering applications.

## Introduction

Protein secretion is the process through which proteins are exported outside of the cell cytoplasm^1,2^. In eukaryotic cells, most proteins destined for secretion are involved in cell-to-cell communication and cell adaptation to external conditions. These proteins are first synthesized as precursor polypeptides at the ribosomes attached to the rough endoplasmic reticulum (ER) and are then co-translationally translocated into the ER lumen, where they undergo folding and post-translational modifications^3–5^ before being directed to the Golgi apparatus. This initial targeting is mediated by an N-terminal signal peptide that directs the nascent polypeptide to the translocation machinery which ensures entrance into the ER lumen^6,7^. The signal peptide is a ∼10-50 aa sequence at the N-terminus of secreted and membrane proteins characterised by a positively charged N-terminus, which facilitates entry into the ER membrane, a hydrophobic core, also required for membrane insertion, and a cleavage site to release the cargo into the ER^6,8–10^.

Secretion plays a key role in the diverse applications of cell engineering. In bioproduction, it simplifies purification of the product by separating it from cell mass and ensures its post-processing through the ER^11–13^. In synthetic biology, as the field evolves from gene circuits to cell circuits, secretion is pivotal for the tuning of intercellular signals^14–16^. In the field of cell therapies, the secretion of cytokines is becoming fundamental to the latest generation of immune cell therapies by boosting their effectiveness and immune activation role^17–19^. However, while in bioproduction the aim is generally to maximise secreted output, for cell therapies, this can cause unwanted side effects such as cytokine storms^19,20^. On the other hand, in synthetic biology, engineering of cell-to-cell communication might require dynamic control over secretion levels^14,16^. The ability to modulate secretion levels of specific target proteins based on needs would thus greatly benefit several research areas in mammalian cell engineering.

Native signal peptides, though optimised by evolution for endogenous protein secretion, were not selected to maximise recombinant protein secretion and exhibit variable performance across cell types, cargos and secretory load. Moreover, endogenous signal peptides have co-evolved with their corresponding protein and are therefore less likely to yield good candidates when transferred across to a heterologous protein^21^. Finally, while computational modelling has so far been useful in predicting natural signal peptide sequences^22–26^, it has not been applied to the generation of novel mammalian signal peptides, leaving a need in the field for identification and screening of a wider array of peptides that can support cell engineering applications.

Historically, signal peptide selection for secretion optimisation has relied on empirical screening or rule-based heuristics^27–29^, yet these methods are inherently limited by the vast combinatorial nature of sequence space. For example, an average mammalian signal peptide of 20-22 amino acids has a sequence space of ∼10^12–13^ possible combinations. While several computational tools exist to predict the presence and cleavage sites of signal peptides (e.g., SignalP^23^, TSignal^22^), their performance can be highly cell-type and protein context-dependent and often does not generalize well to *de novo* prediction. Wu *et al.* ^30^ demonstrated that transformers, attention-based deep-learning models, when conditioned on functional and taxonomic tags, can generate signal peptides with biologically relevant features and potential secretion capabilities^30^. Instead, Teufel *et al*. used pre-trained transformers to estimate quantitative effects on secretion from signal peptides based on the perplexity metric generated by the pre-trained model. Although this study did not involve *de novo* generation, it is a first effort towards quantitative prediction applicable across signal peptide classes^31^. The application of these deep generative strategies capitalises on their ability to capture long-range residue interactions and context-dependent sequence patterns, thereby efficiently modelling the intricate sequence-function relationships that underpin subcellular targeting^32^. However, their application to *de* novo generation has been so far limited to gram positive bacteria^30^.

Here, we fine-tuned a pre-trained autoregressive transformer model on ∼6,000 mammalian signal peptides to make STAMPS (Signal-peptide Transformer for Augmenting Mammalian Protein Secretion). We first used the model to generate 10 synthetic signal peptides and identified 6 that successfully lead to secretion of two different proteins, EGFP and hEPO, and confirmed results are consistent in two mammalian cell lines, HEK293T and CHO-S. We then scaled the testing up to 89 new synthetic signal peptides to gain insights on sequence-to-function relationships. 55 of these led to protein secretion, 44 to a level comparable or higher than the widely adopted Albumin signal peptide, and 10 to a level comparable or higher than IgKL, the IgG kappa light chain leader sequence. Computational analysis of this dataset highlighted the importance of a long hydrophobic core and tightly defined cleavage site to drive high levels of secretion in heterologous protein expression.

Finally, to highlight relevance for therapeutic biomanufacturing, we tested 88 synthetic signal peptides on a VHH-Fc^33^ and compared their secretion yields to an internal benchmark when tested in the industrial CHO G22 cell line within an industrial production workflow. This led to the identification of signal peptides that improve secretion output of a 10-day culture up to ∼2.3-fold. We again confirmed portability across cell lines. These results suggest that STAMPs-generated signal peptides have a degree of transferability across proteins and between CHO and HEK cells. To our knowledge, this is among the first demonstrations that deep-learning-generated signal peptides can successfully drive secretion in a mammalian system, complementing prior studies in bacteria. Our model allows rapid and efficient screening of novel signal peptides offering sequence diversity close to native sequences, a high hit rate of sequences inducing secretion validated *in vitro*, and a varied secretion profile to tune protein secretion for valuable downstream applications. Overall, this work demonstrates how the combination of Transformer-based generative models and oligo pool-based cloning offers a practical, fast and affordable route to optimise signal peptides for mammalian secretion and contributes to widening the toolbox of signal peptides available to cell engineering applications, also supporting the development of more efficient expression systems for biomanufacturing.

## Results

### Data collection and model training

To generate *de novo* mammalian signal peptides, we started by developing and fine-tuning a generative machine learning model. Taking inspiration from previous work by the Arnauld and Nielsen labs^30,31^, we began by filtering Uniprot^34^ for proteins bearing a signal peptide and deriving from mammalian hosts, also considering if secretion had been computationally predicted and/or experimentally validated. The Uniprot database was accessed on 12/09/2023 to download ∼6k mammalian protein sequences with supporting experimental evidence of their secretion and ∼50k protein sequences with potential to be secreted based on sequence similarity (Figure 1A).

**Figure 1.**
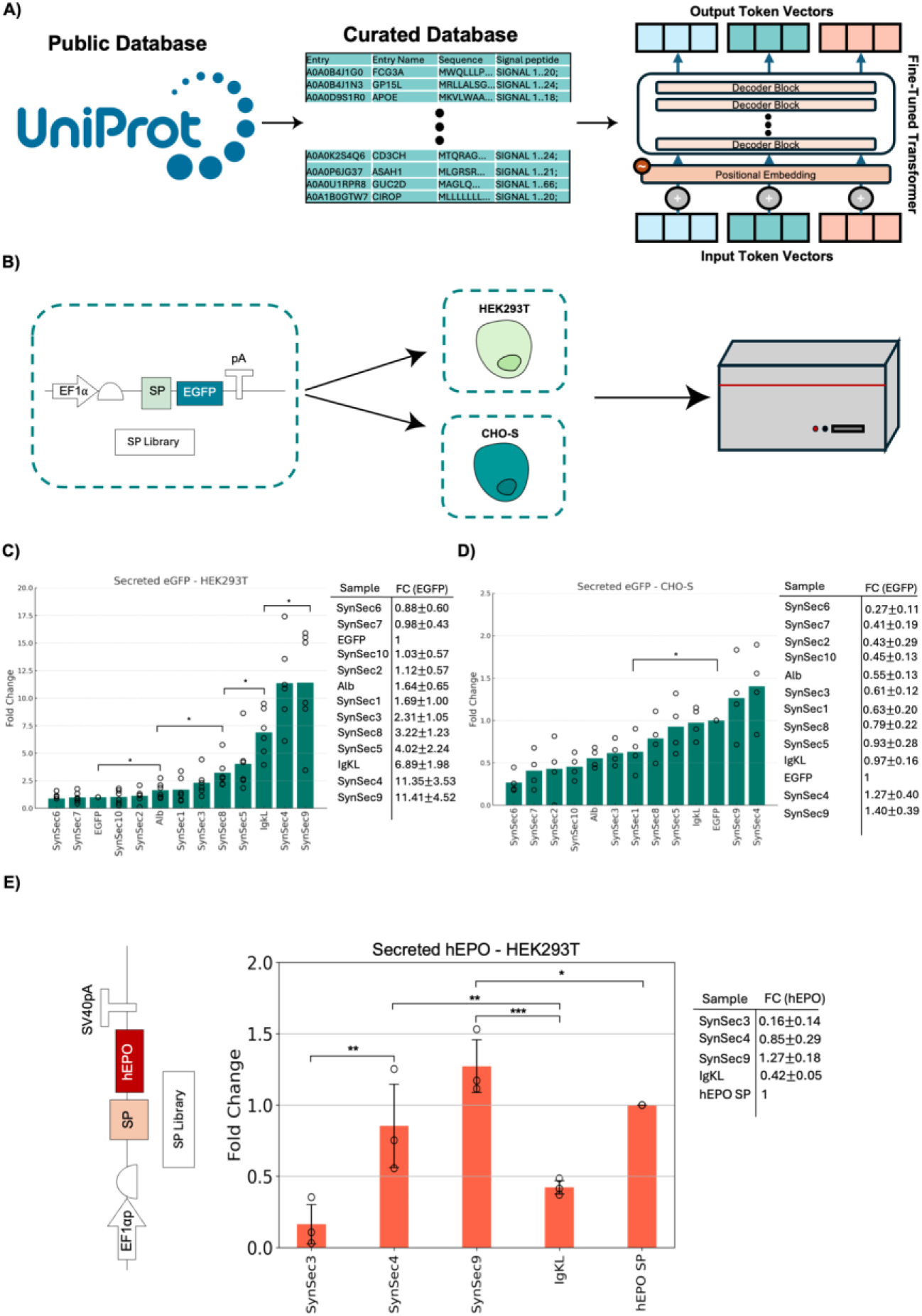
Design and validation of STAMPS. A) Flow chart summarizing data collection, curation, and finetuning of an autoregressive Transformer model. B) Diagram of the experimental workflow from genetic design to host and readout. C) Plate reader quantification of EGFP in the supernatant of transfected HEK293T cells. The data are presented as fold change, calculated by subtracting the signal from untransfected cells in media (blank) from all measurements and then dividing by the fluorescence value of cytosolic EGFP in the supernatant of transfected cells. Each bar represents the mean of three independent experiments, each with two biological replicates. D) Plate reader quantification of EGFP in the supernatant of transfected CHO-S cells. The data are presented as fold change, calculated by subtracting the signal from untransfected cells in media (blank) from all measurements and then dividing by the fluorescence value of cytosolic EGFP in the supernatant of transfected cells. Each bar represents the mean of three independent experiments, each with two biological replicates. E) Diagram of the experimental setup from genetic design to host. E) Bar plot of the mean fold change in hEPO concentration in the HEK293T supernatant for the respective samples. Foldchange values represent the mean ± s.d. from three independent experiments, each with two biological replicates. Significance was assessed in all figures with two-tailed Student’s t-tests (*P < 0.05; **P < 0.005; ***P < 0.0005; ****P < 0.00005; *****P < 0.000005).

Sequences with >99% identity to other entries in the database were filtered out to avoid biasing the model towards specific designs that have many close homologues or duplicates in the database. Additionally, sequences with fewer than 50 amino acids downstream of the signal peptide were also removed to keep the size of the sequences used in model training consistent. 50 amino acids were chosen as a cutoff as it would include the area surrounding the cleavage site and a substantial part of the N-terminus, which are known to influence the efficacy of the signal peptide. After filtering, the portion of the sequence containing the signal peptide element together with 50 downstream amino acids were adopted for model training. The amino acid sequences were tokenised and used to fine tune a Generative Pre-trained Transformer 2 (GPT2) model^35^ (∼85M parameters) resulting in STAMPS (Figure 1A). We validated our autoregressive transformer for mammalian signal peptides *in silico* by classifying the generated sequences with SignalP6.0, the state-of-the-art model for signal peptide classification, and assessing accuracy as the percentage of sequences classified as signal peptides. Then, sequence diversity was assessed via position-wise Shannon entropy over the length of the signal peptide, highlighting a similar level of sequence diversity to naturally occurring sequences (Figure S1).

### *In vitro* validation of model-generated synthetic signal peptides

To validate the generated library of synthetic signal peptides experimentally, we started by selecting as positive control the signal peptide IgKL from the immunoglobulin kappa light chain as positive control. IgKL is commonly used in industry for IgG production and known to lead to high secretion levels. The IgKL signal peptide was cloned downstream of a human EF1α constitutive promoter, a Kozak sequence, and before an enhanced green fluorescent protein (EGFP) for sustained constitutive expression in mammalian cells. Plasmids bearing the EGFP protein either untagged or tagged with the IgKL signal peptide were used to express EGFP in HEK293T cells.

Brefeldin A (brefA) is a mycotoxin known to prevent protein trafficking from the ER to the extracellular space in mammalian cells, thus inhibiting protein secretion. We therefore hypothesised that it could be used to verify whether our secretion control IgKL induces secretion of EGFP by observing an increase in intracellular EGFP upon inhibition of the secretion pathway. To assess the effect of brefA, we transfected HEK293T cells with plasmids expressing EGFP either tagged or untagged with IgKL and either exposed to brefA or not. 30hrs after transfection, we assessed intracellular fluorescence by fluorescence microscopy. Microscopy revealed that brefA exposure differentially affected intracellular EGFP depending on whether it had been tagged with IgKL or not. The increase in intracellular EGFP in the presence of brefA was observed only in the tagged condition, confirming that IgKL acts as a peptide sequence inducing protein secretion (Figure S2).

Once confirmed that our secretion control led to EGFP secretion, we assembled a small library of constructs where ten of the synthetic signal peptides generated by the model were fused to the N-terminus of EGFP (Figure 1B).

These were randomly generated and predicted to lead to secretion of EGFP by SignalP6.0. The selected peptides were cloned in the same expression cassette previously adopted for IgKL testing and constructs were transfected in HEK293T and CHO-S cells via lipofection. To complement our positive secretion control, IgKL, we added a low secretion control already widely validated and adopted for mammalian protein secretion, the albumin signal peptide (Alb). Furthermore, we tested an untagged cytosolic EGFP as a negative control of protein secretion. After 30hrs of expression, the supernatant was harvested, and green fluorescence was measured with a plate reader.

Six out of the ten signal peptides generated by the model (Synsec1, 3, 4, 5, 8, 9) displayed higher levels of secretion compared to our lower secretion control Alb in HEK293T cells, with Synsec1, Synsec3, Synsec4, Synsec8, Syncsec5 and Syncsec9 yielding ∼1x, ∼1.2x, ∼1.5x, ∼2x, ∼6x fold increase, respectively. Synsec4 and Synsec9, displayed ∼1.6-fold increase in secretion compared to IgKL, our high secretion control (Figure 1C).

To investigate the context-dependency of our synthetic signal peptides and their potential transferability between cell lines^36,37^, we performed further validation in CHO-S cells (Figure 1D). Similarly to the results obtained for HEK293T cells, Synsec1, 3, 4, 5, 8 and 9 displayed up to ∼3-fold increase in secretion compared to Alb. Synsec4 and Synsec9 yielded up to ∼1.2-fold increase in secretion levels compared to IgKL. Interestingly, the hierarchy of secretion amongst the same set was remarkably similar in the two cell lines. Surprisingly, we also found that the untagged EGFP control was present in the culture supernatant at high levels for CHO-S cells. This was not expected, given the absence of a signal peptide on the protein. While cell death could be a cause of accidental release of intracellular EGFP in the medium, thus leading to fluorescence in the supernatant, we had designed the experiment to avoid cell death by adopting 20,000 cells per well at seeding stage and limiting expression to 30hrs.

Heterologous expression of proteins in mammalian cells is known to impose a burden to the host resulting in growth deficits which can in turn affect product quality and yield^12,38–40^. To investigate the effects of synthetic signal peptides on cell growth and confirm that the difference in protein secretion was not due to differences in cell growth performance, we used an automated cell counter to confirm the live cell counts 30hrs after expression of tagged or untagged EGFP. The only change in viability detected was for SynSec9 though the total number of live cells after 30hrs of expression was not significantly different from other signal peptides (Figure S3).

### Secretion of hEPO with deep learning-generated signal peptides

Signal peptides have been shown to have a high degree of protein specificity^29,41^. While there is no definitive evidence of the mechanistic reasons for such specificity in the literature, several studies point towards both the sequence and the secondary structure formed by the cleavage site and the protein’s N-terminus, which can either promote, dampen or inhibit secretion^42–44^.

To test the transferability of our newly identified secretion peptides we selected a medically relevant protein, the human erythropoietin (hEPO), and designed a new set of constructs where Synsec3,4 and 9 were cloned upstream of the hEPO gene and were expressed in HEK293T cells. hEPO is a human hormone produced in the kidneys and signals the bone marrow to produce red blood cells by differentiating progenitor cells^45^. Recombinant hEPO is a widely used drug for cases of anaemia associated with multiple clinical conditions. Currently, the most common forms of recombinant hEPO are produced in CHO cells^46^. However, the non-human glycosylation patterns from CHO cells have been shown to potentially lead to undesired immune responses^47–49^. Consequently, recent studies have tried to adapt the production of hEPO to HEK293 cells. Nonetheless, secretion from HEK293 cells is normally much lower than CHO cells^50^.

Synsec3, 4 and 9 were selected alongside the IgKL control and hEPO’s native signal peptide to include two high secretors, Synsec4 and Synsec9, one low, Synsec3, a high control, IgKL, together with the protein’s cognate signal peptide sequence (Figure 1E). To quantify hEPO levels, we performed an ELISA on cell culture supernatant. After thirty hours of expression, we immobilized hEPO on an antibody pre-coated plate, bound it with a biotinylated detection antibody and performed the ELISA with streptavidin-HRP (horseradish peroxidase). The hierarchy of secretion of the tested peptides with respect to each other and the controls remained unchanged in the shift from EGFP to hEPO. We also found that Synsec4 and Synsec9 yielded up to ∼0.8-fold and ∼1.3-fold increase in secretion compared to the native signal peptide and ∼2-fold and ∼3-fold increase when compared to IgKL, respectively (Figure 1E).

### High throughput testing of synthetic mammalian secretion peptides on a fluorescent protein

Initial validation of our autoregressive transformer model confirmed its ability to generate signal peptides that lead to different levels of secretion of a test protein, i.e. EGFP and hEPO. We then reasoned that if we could generate a larger library of candidates, this would give us the opportunity to explore the efficiency variability of our model-generated candidates but also to investigate the relation between amino acid composition and secretion efficiency with the possibility to identify design rules for “optimised” secretion peptides.

We thus scaled up the throughput of our testing by nearly an order of magnitude. Similarly to the initial validation, we adopted SignalP6.0 filtering and selected 96 signal peptides. These were then cloned in the same EGFP expression construct used for the preliminary validation but this time adopting an oligo pool-based approach^51^. Briefly, following the work by Subramanian *et al*.^52^, each signal peptide was tagged on either side with orthogonal primer sequences^46,47,50,51,52^ containing typeII restriction enzyme sites. Orthogonal amplification of each signal peptide sequence allowed the demultiplexing of each oligo from the pool followed by cloning into our backbone via Golden Gate cloning (Figure 2A, B, C). 89 out of 96 total signal peptides were successfully cloned and were then tested in HEK293T cells following the same protocol as the one adopted for the initial model validation. We retained EGFP, Alb and IgKL as our controls for no, low and high secretion, respectively. Out of the total, 34 peptides led to secretion levels higher than Alb but lower than our high secretion control IgKL (∼38%), while 10 signal peptides led to secretion levels higher than IgKL (∼11%) (Figure 2D).

**Figure 2.**
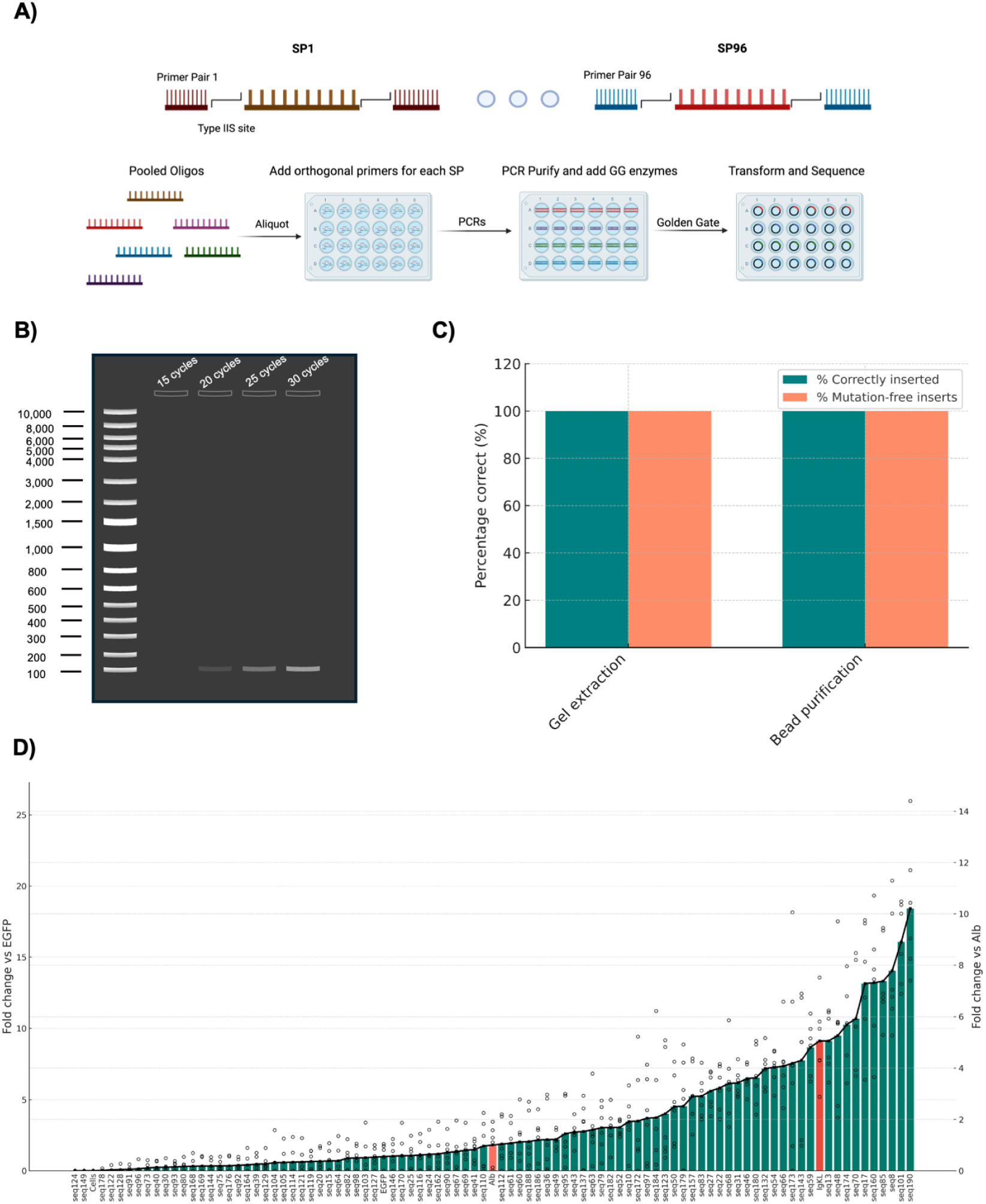
High-throughput screening of synthetic signal peptides from pools of oligonucleotides. A) Diagram representing the experimental setup for cloning of signal peptides from pools of oligonucleotides via orthogonal amplification and Golden Gate cloning. B) Representative image of a gel electrophoresis showcasing different cycling conditions for orthogonal amplification of specific oligonucleotides from a pool. C) Testing of purification via gel extraction or magnetic beads-based purification measured by the number of correct colonies after transformation of the cloned pool-amplified oligonucleotides. D) EGFP fluorescence in the supernatant of transfected HEK293T cells. Values are plotted as fold change with respect to cytosolic EGFP (left) and Alb-EGFP (right). Sample fluorescence was corrected by subtracting the signal from untransfected cells in media (blank), then normalized by dividing by the fluorescence of cytosolic EGFP or Alb-EGFP measured in the supernatant of transfected cells. Each bar represents the mean of three independent experiments, each with two biological replicates.

### Sequence-to-function mapping reveals importance of the leucine core and a preserved signal peptide cleavage site

The field of protein secretion has made significant strides in understanding the sequence determinants of signal peptides, however, it still lacks a quantitative understanding of how amino acid changes can impact secreted output. Even though it is limited to a single protein and cell line, the data obtained from the higher throughput screen of signal peptides allowed us to investigate the relation between the amino acid composition of the signal peptides and their effect on the secretion of a commonly used reporter, EGFP.

To do so, we analysed our semi-quantitative results by mapping sequence composition and features to secreted output. We began by splitting up the fold-change data using a k-means clustering algorithm which led to 4 classes of secretion outputs (class 1 to 4, with 1 being the lowest and 4 the highest). Upon further examination, we merged class 3 and 4 due to data scarcity in the very high secretor class, class 4 (Figure S4). We then looked at amino acid abundance at each position of the sequence for the whole dataset and then split it by secretion class. The only notable differences between secretion classes 2 and 3 compared to class 1 were found in the abundance of leucine in the central region of the signal peptide and the overrepresentation of alanine at the −1 position in class 3 (Figure S4).

Furthermore, we analysed the signal peptide cleavage site position and how it changes based on the secretion class to which the respective sequences belong. The cleavage site position was computationally predicted with the SignalP6.0 model^23^. All the signal peptides were 20 amino acids long, therefore, the expected cleavage position was 19-20, as the sequences are zero indexed. We observed a stark difference in the secretion classes cleavage site position. The higher the secretion levels the more the distribution of cleavage site positions centres on position 19-20. In fact, more than half of the non-secreting signal peptides had a predicted cleavage position at 23-24 (Figure S4).

### High throughput synthetic signal peptides improve VHH-Fc production in industrial CHO G22 cells

After confirming that our model reliably generates high-performing mammalian signal peptides, beyond the initial screen of 10 sequences, we decided to apply a large set of synthetic signal peptides to a therapeutic VHH-Fc, cAbBcII10-Fc^33^. We selected 88 signal peptides from our generation run comprising 200 sequences used for the test with EGFP and hEPO. These were chosen after filtering for likelihood of constituting a eukaryotic signal peptide in front of cAbBcII10-Fc with SignalP 6.0. The signal peptides were cloned via Golden Gate after the start codon of the VHH-Fc in a plasmid vector for transient expression. These were then transfected in the CHO G22 industrial cell line and cultured for 10 days post-transfection following an internal industrial manufacturing workflow before quantification via biolayer interferometry (Figure 3A). The raw protein concentration data inferred from biolayer interferometry appeared highly variable (Figure S5). However, via winsorization of the mean (see Methods section), we found that 54 out of 88 signal peptides led to VHH-Fc secretion levels comparable to or higher than the AstraZeneca internal benchmark (Figure 3B). We also performed a quartile analysis where we quantified the frequency with which each sample’s repeats would appear in the top quartile of the aggregated data (see Methods section).

**Figure 3.**
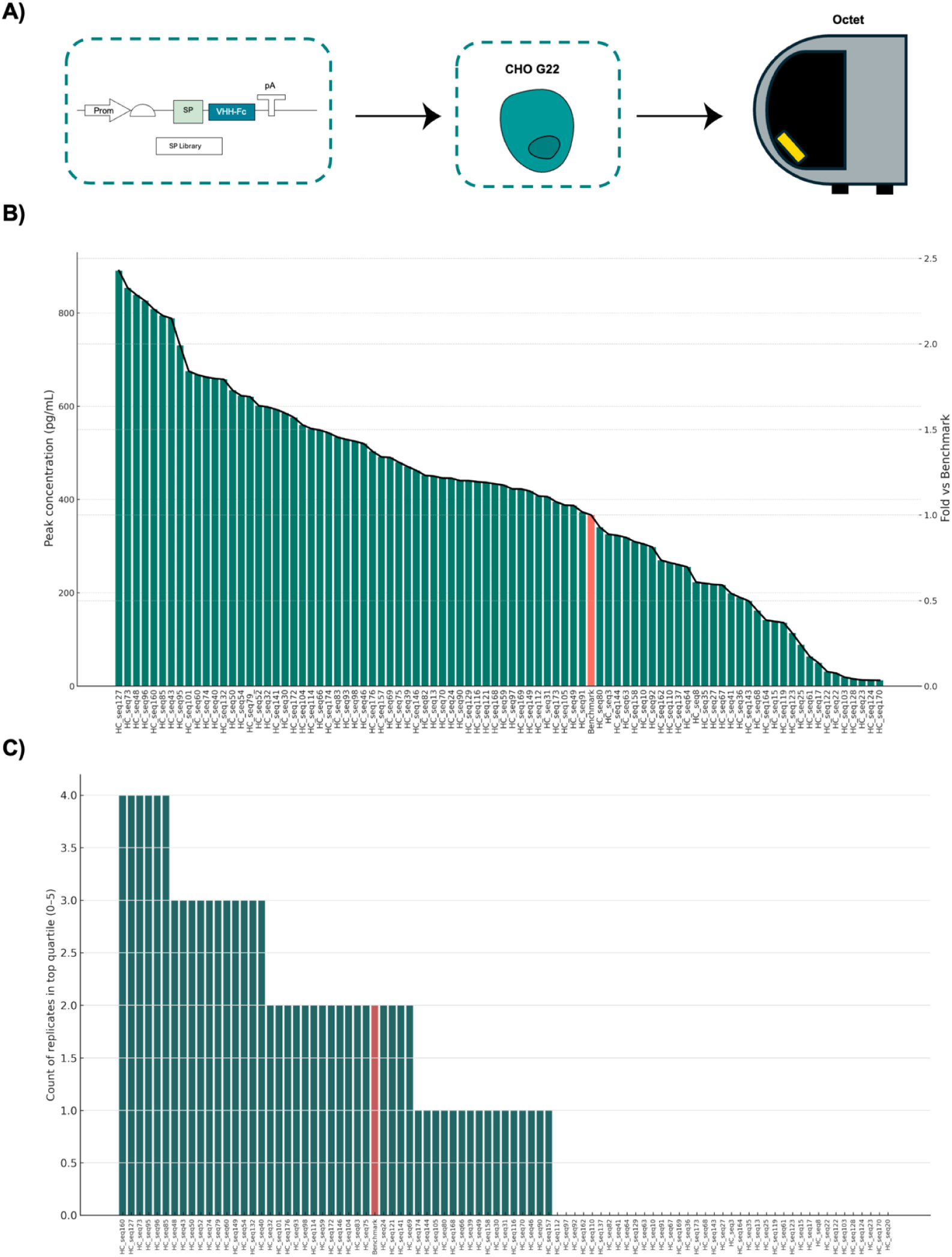
Synthetic signal peptides lead to improvements in VHH-Fc secretion in CHO cells. A) Diagram representing the experimental setup from genetic construct to host and measurement modality. B) Concentrations calculated from biolayer interferometry binding kinetics measurements from a Sartorius Octet of a humanised VHH-Fc from crude CHO G22 supernatant after 10 days of expression in deep 96 well plates. (left) Each bar represents the 90% winsorized mean of five technical repeats from four independent experiments. (right) Fold change with respect to the industrial benchmark. C) Quartile counts based on the top quartile of the aggregated data from 4 independent experiments comprising of 5 technical repeats. Each bar represents the mean of four independent experiments, three with one biological replicate and one with two.

Through the quartile analysis, 37 signal peptides appeared at the same or higher frequency in the top quartile when compared to the benchmark (Figure 3C). Furthermore, 6 signal peptides appeared twice as frequently in the top quartile as the benchmark, and all of these were also in the first 8 signal peptides according to the winsorized means with improvements in supernatant VHH-Fc concentration of up to ∼2.3-fold. Finally, comparing the VHH-Fc with the previous EGFP datasets highlighted the portability of some of our synthetic signal peptides. For example, from the top 10 sequences for secretion output from the EGFP test (seq59, seq143, seq174, seq70, seq17, seq160, seq85, seq8, seq101, seq190), 9 out of 10 are also included in the BLI dataset. Of those 9, 4 sequences (seq160, seq85, seq48 and seq101) are found in the top 10 secretors on the VHH-Fc. Notably, the EGFP test was carried out in HEK293T cells, while the VHH-Fc secretion experiment in CHO G22 cells, suggesting that these signal peptides exhibited not only consistency amongst different proteins but also amongst expression hosts.

## Discussion

Signal peptides (SPs) are short N-terminal sequences that guide nascent polypeptides to the endoplasmic reticulum and thereby constitute a rate-limiting step in recombinant protein production^41,42^. Swapping or redesigning the SP is one of the levers that can be pulled to raise secretion titre directly. Despite their strategic value, only a handful of mammalian signal peptides have been characterised in depth to date, and systematic efforts to build novel variants remain sporadic^29^. Herein, we report how a transformer model trained on known SPs and combined with routine bioinformatics filtering can propose synthetic sequences that match, or exceed, the performance of established signal peptides in professional mammalian cell secretors.

Earlier secretion improvements in the literature have relied almost entirely on empirical replacement. Substituting an antibody’s endogenous leader with the signal sequence from human serum albumin or azurocidin raised titres 1.5- to 2-fold in CHO cells, an observation first reported by Kober and co-workers^53^. Broader panels have been screened in bacterial hosts. Brockmeier *et al.* catalogued natural SPs in *Bacillus subtilis* and documented order-of-magnitude shifts in extracellular enzyme levels between otherwise identical constructs^21^. Yet such empirical efforts do not scale well as a 25-residue leader spans roughly 10^32^ possible sequence variants, far beyond what can be covered experimentally.

Algorithms such as SignalP^23^ and TSignal^22^ can identify whether a sequence is an SP and map the likely cleavage site with high accuracy, but they do not rank signal peptides by secretion strength. To address that gap, O’Neill *et al.* trained a regression model on thousands of SP-protein pairs, screened ∼40,000 candidates in silico and identified variants that doubled production of a single-chain antibody fragment. The model’s predictive power remained modest (R² ≈ 0.65) and weakened for full antibodies, highlighting the context dependence of SP efficacy^29^. Generative approaches attempt to bypass those constraints by generating new sequences directly. Wu *et al.* demonstrated that an autoregressive Transformer trained on bacterial SPs could produce signal peptides that performed on par with top industrial standards when fused to secreted enzymes in *Bacillus*^30^.

Extending this concept to mammalian expression, in this work we fine-tuned an autoregressive transformer model on mammalian signal peptide sequences to generate synthetic ones with ∼85% *in silico* and ∼60% *in vitro* accuracy and high sequence diversity. The synthetic signal peptides were validated first with ten sequences tagging EGFP across two cell lines, HEK293T and CHO-S cells. We then tested and confirmed the same sequences drive secretion of hEPO in HEK293T cells with consistent peptide ranking across proteins. Finally, we performed a larger screen on 89 candidates driving secretion of EGFP in HEK293T. In these experiments, synthetic sequences achieved the highest outputs across conditions but also displayed a broad range of secretion levels.

We analysed our dataset of 89 *in vitro* characterised synthetic signal peptides to obtain insights on amino acid changes with quantitative effects on secretion. Our main observation was that higher hydrophobicity of the signal peptide core positively correlated with increased secreted output and previous studies have shown the importance of the hydrophobic region for ER translocation^54–56^. Finally, we identified that signal peptide cleavage sites positioned downstream of the SP-protein junction would severely hamper secretion levels. While we could not verify whether this was due to reduced secretion efficiency or protein-related effects such as misfolding, this would be consistent with previous studies. Güler-Gane *et al*. have previously observed how changing one or two amino acids around the cleavage site with the same signal peptide and target protein drastically altered secretion levels in HEK293T cells^42^. Extending our study to other proteins and cell lines will also be a beneficial step for future studies. Additionally, our transformer was trained on a heterogeneous set of SPs rather than on protein-specific or organism-specific examples which could prevent the model from reaching global optima.

To demonstrate applicability of our approach to biomanufacturing, we then used a set of 88 signal peptides from the same model output to produce a VHH-Fc in the industrial cell line CHO G22. We were able to identify candidates with up to ∼2.3-fold increase in secretion compared to an industrially relevant benchmark. Moreover, the signal peptides that were present in both HEK293T and CHO G22 cell experiments displayed consistent ranking, suggesting portability of the synthetic sequences. Our validation of the model-generated signal peptides in CHO G22 cells producing a VHH-Fc confirms that deep-learning-guided SP design is a viable strategy for improving secretion in relevant mammalian hosts producing antibody-like proteins.

In the future, we envisage that larger quantitative datasets, cleavage-site classifiers and multi-objective optimisation frameworks will further increase reliability and extend the method’s applicability to a wider range of biotherapeutics. These design pipelines could also incorporate folding or glycosylation predictors alongside secretion models and adopt iterative generate-test-refine cycles, gradually biasing the model toward SP features that benefit each individual protein in the context of its secreting host.

## Methods

### Model fine-tuning

For signal peptide sequence generation, we used the small version (∼80M parameters) of the autoregressive GPT-2 model^35^ fine-tuned for 20 epochs using Pytorch and the HuggingFace Transformer library. A curated set of ∼6,000 SwissProt signal peptides was encoded with an ESM tokeniser. Training, carried out with batching and gradient accumulation steps, minimised token-level cross-entropy using the Adam optimiser (default hyperparameters β1=0.9, β2=0.999, ɛ=10^-8^). Training was stopped once validation loss plateaued following ∼20 epochs. Generation accuracy was measured by classifying the generated sequences with SignalP6.0 as signal peptide or non-signal peptide sequence. Sequence diversity was calculated as the position-wise Shannon entropy of the generated sequences versus the experimentally validated natural mammalian signal peptides from Uniprot.

### Plasmid design and preparation

The first ten signal peptides used to validate our model were assembled by ordering the forward and reverse DNA strands as single-stranded oligonucleotides from IDT, annealed by gradually decreasing temperature from 98°C to room temperature at 0.1°C/s, PCR purified in columns according to the manufacturer’s protocol (2814, Qiagen), and cloned into our expression vector via Golden gate cloning using the EMMA protocol^57^ with Esp3I (R0734, New England Biolabs) and T4 DNA ligase (M0202L, New England Biolabs). Ligations were transformed in chemically competent *E. coli* DH5α by heat shock at 42°C for 30s and 45min outgrowth in 1mL Luria Broth (L3522, Sigma-Aldrich). 100μL were then plated on agar plates supplemented with 100μg/mL ampicillin (A9518, Sigma-Aldrich). The antibiotic-resistant colonies were then inoculated in 5mL of Luria Broth the following day and incubated overnight at 37°C. The culture was then midi prepped (12843, Qiagen) according to the manufacturer’s protocol the following day to extract the plasmid DNA, ready to be diluted for transfection. A complete list of plasmids included in this study can be found in Related File 1. Plasmid maps annotated for specific parts and sequences used in this paper can be found in the Source data file. Primers used in this study can be found in Supplementary Data.

### Mammalian cell culture and transfection

HEK293T cells (HEK293T, ATCC CRL3216) were cultured in T75 flasks (156340, ThermoFisher Scientific) using DMEM (Dulbecco’s modified Eagle Medium) supplemented with 1mM sodium pyruvate, 1x GlutaMax™, 0.4mM phenol red, and 25 mM glucose (10569010, Thermofisher Scientific), along with 10% FBS (A5670701, Gibco). Cells were kept in an incubator at 37 °C and 5% CO_2_ and were passaged upon reaching 70% confluency. 1mL of trypsin-EDTA (0.5%, no phenol red, Gibco) was used for cell detachment during passages, and 1mL of PBS (phosphate buffered saline, Merck) was used for cell washing. CHO-S were cultured in MEMa (M2279, Sigma) supplemented with 10% FBS (A5670701, Gibco), 1% non-essential amino acids (11140050, Thermofisher Scientific), and 1% L-glutamine (25030081, Thermofisher Scientific) in baffled bottom 125mL shake flasks and split every 48hrs (4116-0125, Thermofisher Scientific). HEK293T cells were seeded 1 day before transfection at 60,000 cells/well in 48w plates (10062-898, VWR) for the transient transfections. Details on the transfection mixes can be found in Supplementary Data. The complete mix was incubated for 30min pre-transfection and 2 days after. CHO-S cells were seeded for transfection at 20,000 cells/well in 48w plates one day before transfection to minimise cell death and protein release. Details on the transfection mixes can be found in Related File 2. The complete mix was incubated for 25min at room temperature pre-transfection and 30 hours after.

### VHH-Fc expression

Recombinant proteins were expressed in suspension-adapted CHO G22 cells and maintained in an AstraZeneca proprietary medium supplemented with methionine sulfoximine (MSX) and copper sulfate. Cell cultures were resuspended to a final density of 4×10^6^ viable cells/mL in a pre-warmed fresh media containing 1% (v/v) dimethyl sulfoxide (Merck, D2438) and 1X antibiotic/antimycotic solution (Merck, A5955). Transfection was performed by dispensing 0.75mL of cell suspension into 96-deepwell plates (Eppendorf, 951033502) and adding 0.75µg plasmid DNA complexed with PEI Max® (Polysciences, 24765) at a PEI:DNA ratio of 5:1 (w/w). Cell cultures were incubated at 37°C in a humidified incubator (≥80% relative humidity, 5% CO2) with orbital shaking at 500rpm (25mm diameter). Four hours post-transfection, proprietary feed supplements F-09 and F-10 were added at 5% (v/v) and 0.6% (v/v), respectively, and temperature was shifted to 34°C. Additional feeding was performed on days 3 and 6. The cultures were incubated for 10 days and conditioned medium containing secreted proteins was harvested by centrifugation at 3200rpm for 30min.

### Cloning from oligo pool

Individual oligos were designed by adding orthogonal primer pairs with identical melting temperatures at either end of the oligos as described in Subramanian *et al*.^52^, followed at the 5’-end and preceded at the 3’-end by type IIs restriction sites as described in Lund *et al*.^51^ The designed oligos were ordered as a pool from Twist Biosciences. The orthogonal primers allow selective recovery of individual oligos from the pool via PCR which we carried out with Phusion polymerase according to the manufacturer’s protocol (M0530L, New England Biolabs), by double-stranding and amplification of only one oligo per reaction with at least 30 cycles for enrichment of the target oligo. The PCR reactions were purified either via a bead-based method (A63881, Beckman Coulter) or gel extraction according to the manufacturer’s protocols (28704, Qiagen). We proceeded with the AmpureXP beads-based method as it makes it easier to handle large amounts of samples. The restriction sites allow cutting of only the double-stranded DNA for insertion into a vector backbone via Golden Gate cloning which was performed as described in Martella *et al*.^57^ Transformations in DH5α chemically competent cells were carried out in PCR tubes in a thermocycler for efficient handling of high volumes of samples. Finally, minipreps followed the kit manufacturer’s protocol (27106, Qiagen) but cultures were only 1.5mL in 2mL 96 deep-well plates (780270, Greiner) sealed with a breathable membrane (Z380059, Sigma Aldrich).

### Cloning of signal peptide library for VHH-Fc expression

A library comprising 88 distinct secretion peptides and one control sequence was synthesised by Integrated DNA Technologies (IDT) using CHO cell codon optimisation. Each DNA fragment was flanked by BsaI recognition sites to enable directional assembly into the acceptor vector pBETX02. The signal peptide sequences were cloned upstream of the start codon of a VHH-human-Fc fusion, cAbBcII10-Fc^33^. Golden Gate assemblies were performed using the NEBridge® Golden Gate Assembly Kit (E1601S, New England Biolabs) in a 2µL final volume containing approximately 1ng of DNA insert and 15ng of pBETX02 vector. Reactions were dispensed using an ECHO 525 acoustic liquid handler (Beckman Coulter) and subjected to the following temperature-cycling program: 37 °C for 1 min, 16 °C for 1 min, repeated for 30 cycles, followed by 60 °C for 5 min. Assembly reactions were transformed into Z-competent E. coli cells in 96-well format, which were incubated on ice for 5 min, and plated on 2×TY agar (YDA1L, Formedium) containing carbenicillin (100 µg/mL) (C1613, Sigma Aldrich). Single colonies were inoculated into 96-deep-well plates containing 1.2 mL 2×TY + ampicillin (100 µg/mL) and cultured at 30 °C, 280 rpm for 16hrs. A 20µL aliquot of each culture was used to initiate a secondary 96-deep-well plates culture under the following conditions, 16hrs at 37°C, 320 rpm. Cells were harvested by centrifugation, and plasmid DNA was purified using the QIAprep 96 Plus Miniprep Kit (27291, Qiagen). DNA concentration and purity were assessed using a Stunner (Unchained Labs) spectrophotometer. Sequence-verified constructs were confirmed by Sanger sequencing, aligned to the reference VHH-Fc design, and normalised to 30ng/µL for transfection.

### Fluorescence, ELISA and cell count assays

Measurements of protein secretion were carried out via plate reader measurement of the supernatant of secreting cells. Thirty hours after transfection, the supernatant was aspirated from individual wells and spun down at 200×*g* for 5min in 1.5mL Eppendorf tubes to get rid of floating cells. 100µL of supernatant were then transferred to black bottom 96-well plates (3915, Corning) for plate reader acquisition in a Tecan Spark (Tecan). mKate fluorescence was measured with a 550nm excitation wavelength and a 600nm emission wavelength with optimal gain. EGFP fluorescence was measured with a 485nm excitation wavelength and 535nm emission wavelength and optimal gain. For live cell counting experiments, transfected cells in 48w plates were washed in DPBS detached in 100µL of 1% trypsin-EDTA, and 200µL of DMEM + 10% FBS was added to a total volume of 300µL. Lastly, 2.5 µL of solution 18 (910-3018, Chemometec) was added to 50µL of cell suspension and live cells were counted using the Nucleocounter (Nucleocounter NC250, Chemometec). Cell counts can be found in the Source Data File. The hEPO ELISA assay was also carried out 30 hours after transfection in HEK293Ts described above, and the assay was done following the manufacturer’s protocol and exclusively using the kit’s provided reagents (BMS2035-2, Thermofisher Scientific).

### VHH-Fc quantification via biolayer interferometry

Crude antibody quantification in clarified culture supernatants was performed using an Octet® RH16 biolayer interferometry (BLI) system (Sartorius) equipped with Octet® Protein A (ProA) Biosensors (Sartorius, 18-5010). A reference antibody of known concentration was serially diluted in fresh culture medium to generate a standard curve. Sample preparation involved diluting clarified supernatants 1:5 (v/v) in fresh culture medium to ensure measurements fall within the linear range. The BLI assay was run using standard ‘Basic Quantitation with Regeneration Experiment – Protein A, G or L biosensors_8Channel_96well plate’ settings. Data analysis was conducted using Octet® BLI Discovery and Analysis Studio Software, with antibody concentrations determined by interpolation from the standard curve.

### Microscopy

Fluorescence imaging was performed on a Nikon ECLIPSE Ti2 with standard filter cubes for EBFP/CFP, ∼390/40nm excitation wavelength (EX), 460/50nm emission wavelength (EM), EGFP∼470/40nm EX, 525/50nm EM; cells were imaged in tissue-culture–treated 48w plates in PBS or full medium.

### Statistics

All statistical analyses were performed on biological replicate measurements unless otherwise specified. For all pairwise comparisons between conditions, statistical significance was assessed using a two-sided paired Student’s t-test. To limit the influence of extreme values on summary statistics, means were calculated as 10% winsorized means: within each condition, the lowest and highest 10% of replicate values were replaced by the values at the 10th and 90th percentiles (Figure 3), respectively, prior to computing the mean. Exact sample sizes and P values are reported in the corresponding figure legends, and differences were considered statistically significant at P < 0.05.

### Data Availability Statement

All source data and plasmid maps for the data sets presented in the manuscript are available with this publication as related manuscript files. Constructs are available upon request.

### Software availability statement

The model is deposited on HuggingFace comprising of training and model details, metrics and weights at the following address: Jacopo-gab/230912GPT2_fine_tuned_SP_GPT2_config_ESM_tokenizer_6kSwissProt_20epochs.

## Acknowledgements

The authors acknowledge the support of Astra Zeneca and the EPSRC Centre for Doctoral Training in BioDesign Engineering (EP/S022856/1) (to J.G., C.K. and F.C.). FC and CK were also partly funded by the Bezos Earth Fund through the Bezos Centre for Sustainable Protein (BCSP/IC/001) and the UK National Alternative Proteins Innovation Centre (NAPIC), which is an Innovation and Knowledge Centre funded by the Biotechnology and Biological Sciences Research Council (BBSRC) and Innovate UK (BB/Z516119/1). FC is supported in part by the Engineering and Physical Sciences Research Council under the EEBio Programme Grant (EP/Y014073/1) and by the Chan Zuckerberg Initiative.

## Competing interests

The authors declare no competing interests.

## Notes

### Competing Interest Statement

The authors have declared no competing interest.

