## Supplementary Information for "STAMPS: Signal-peptide Transformer for Augmenting Mammalian Protein Secretion"

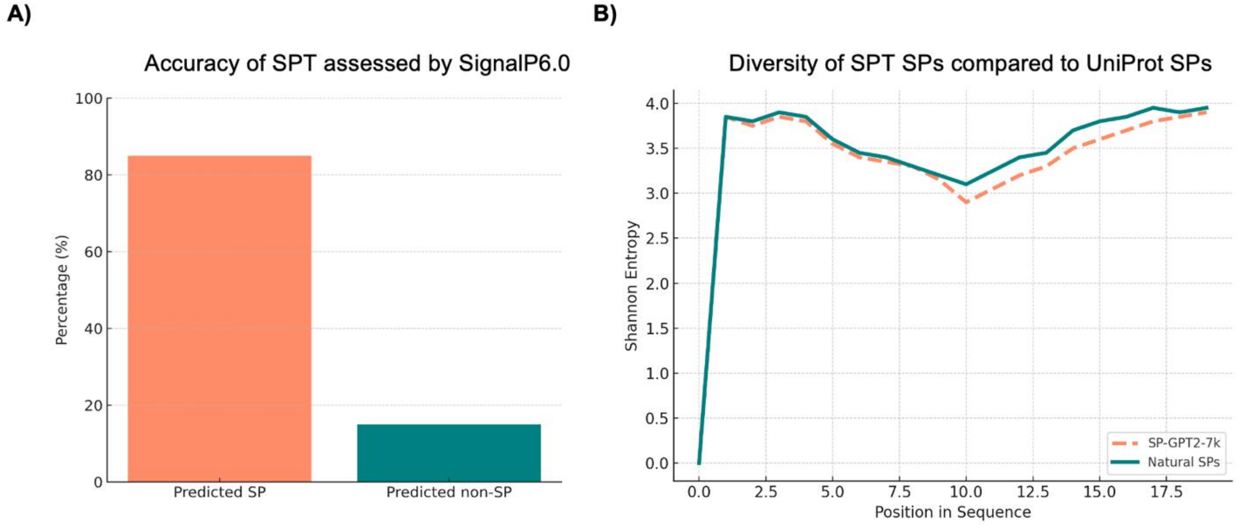


**Figure S1. STAMPS is accurate at generating diverse signal peptide sequences.** A) Accuracy of STAMPS as assessed by whether the generated sequences are classified as eukaryotic signal peptides or not by SignalP6.0 (n=200). B) Diversity of the amino acid sequence of STAMPS generated signal peptides measured using Shannon entropy and compared to natural sequences from UniProt.

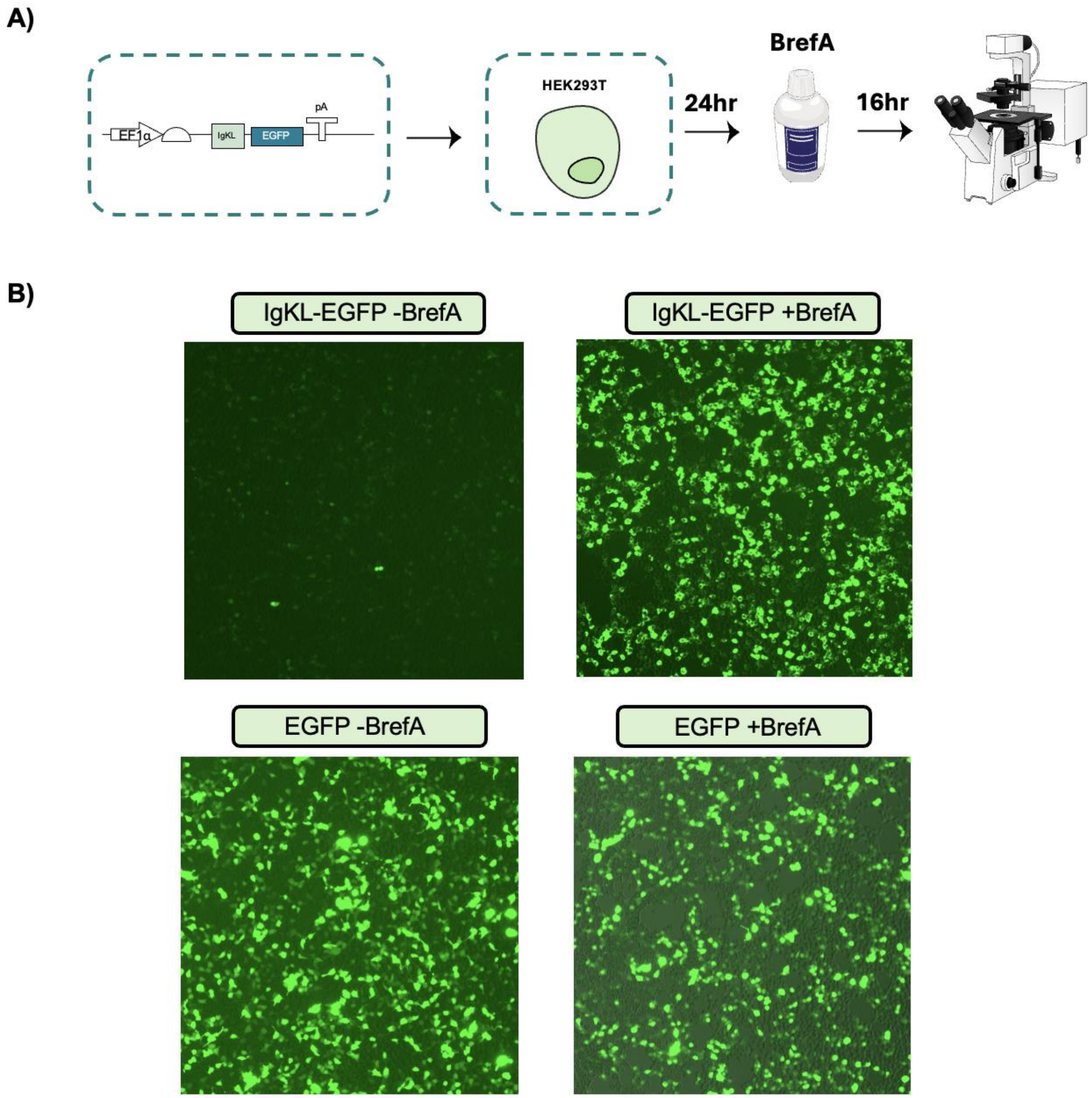


**Figure S2. IgKL induces EGFP secretion in HEK293T cells which is inhibited by Brefeldin A.** A) Diagram illustrating experimental setup from construct design to transfection, exposure to BrefA and visualization. B) Representative fluorescence microscopy images with a fluorescence green channel overlaid to its corresponding brightfield of HEK293T cells transfected with constitutively expressed EGFP either tagged with IgKL or untagged and exposed or unexposed to BrefA (n=2).

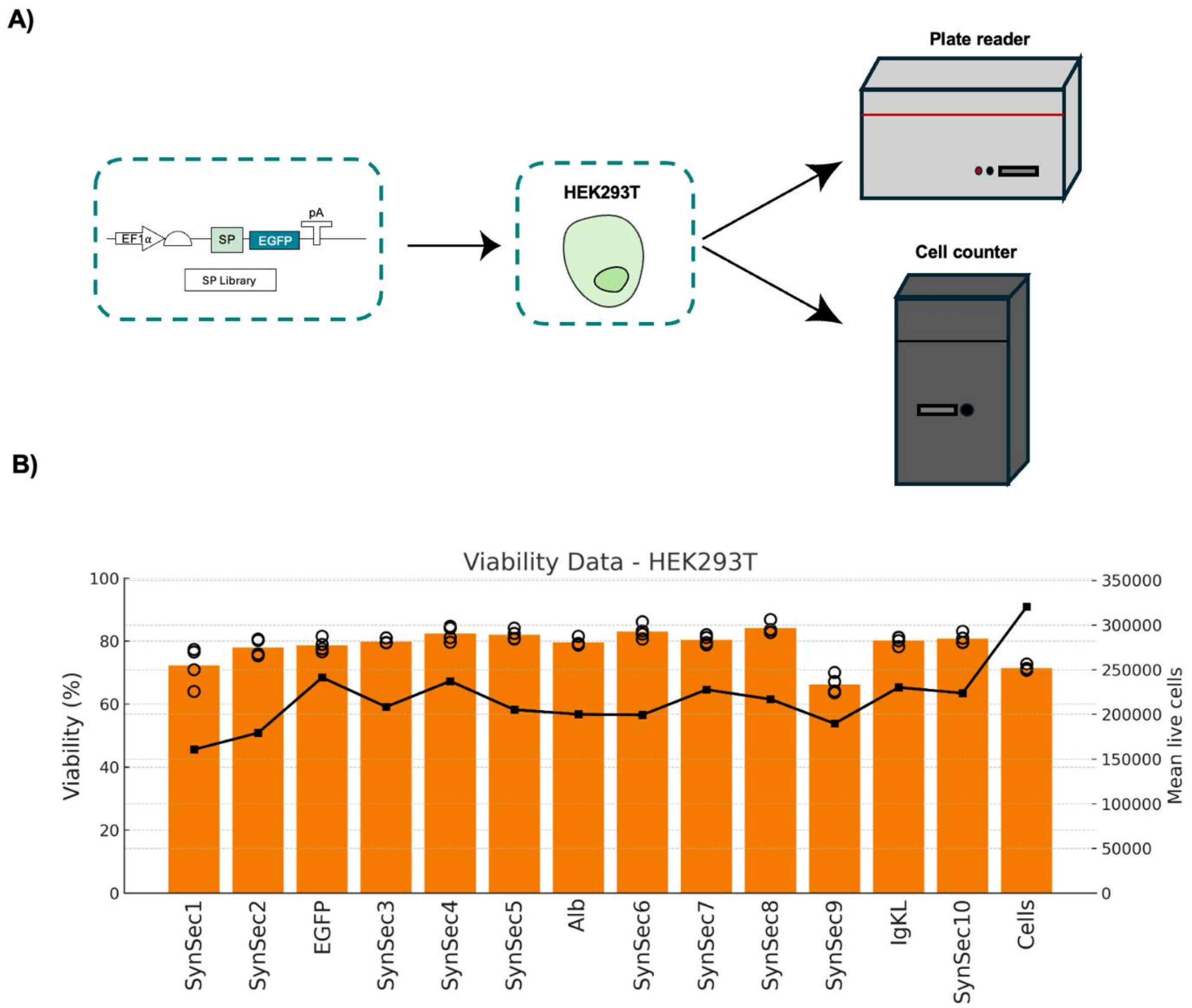


**Figure S3. STAMPS synthetic signal peptides do not significantly impair cell viability after 30hrs of expression.** A) Diagram illustrating experimental setup from construct design to transfection, cell counting and fluorescence measurement. B) Bar chart displaying the viability and mean live cells per well of HEK293Ts after 30hrs of expression of EGFP either secreted with controls or our synthetic signal peptides, cytoplasmic or transfected with an empty plasmid (n=2).

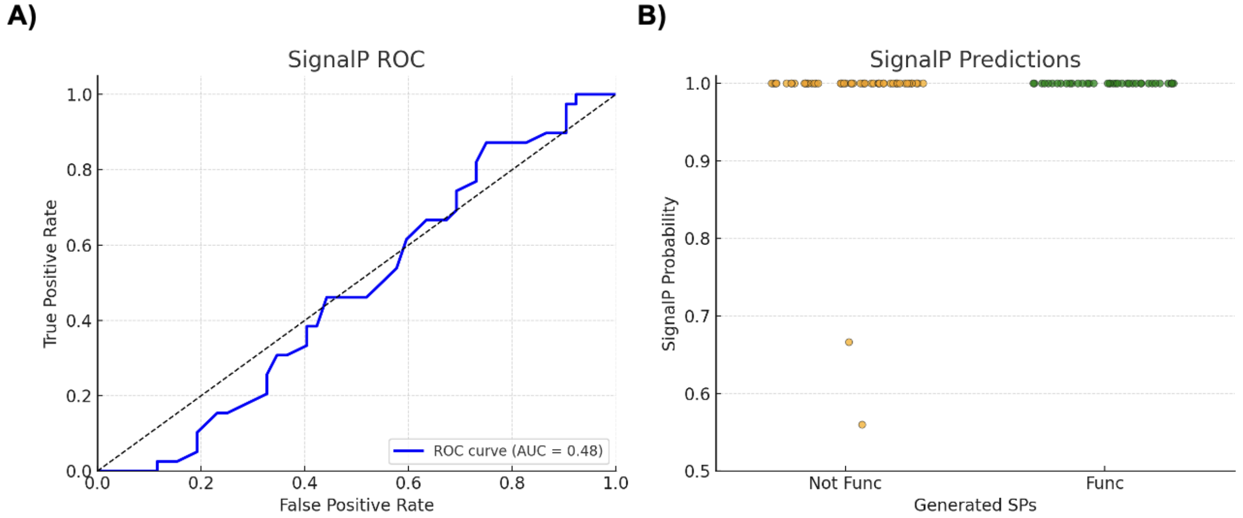


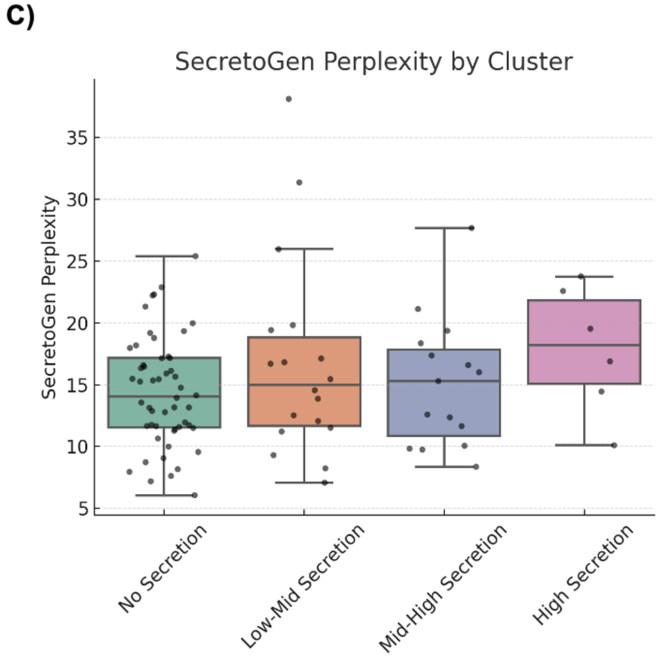


D)

| Panel | Description | Key Result |
| --- | --- | --- |
| A | SignalP ROC curve | AUC = 0.48 |
| B | SignalP prediction probabilities | TP = 50% FP = 50% |
| C | SecretoGen perplexity by secretion cluster | No Secretion – High Secretion (p-value = 0.10) |

**Figure S4. Benchmarking signal peptide models with our dataset.** A) ROC of the SignalP6.0 predictions on our synthetic signal peptides validated in HEK293Ts as either inducing or not inducing secretion. B) Dot plot comparing the probability of being a signal peptide assigned by SignalP6.0 to our synthetic signal peptides across the ones found to be functional or non-functional in HEK293Ts. C) The perplexity score assigned by Secretogen to our signal peptides grouped via k-mean clustering into four classes based on secretion intensity. D) Summary table of the benchmarks displayed.

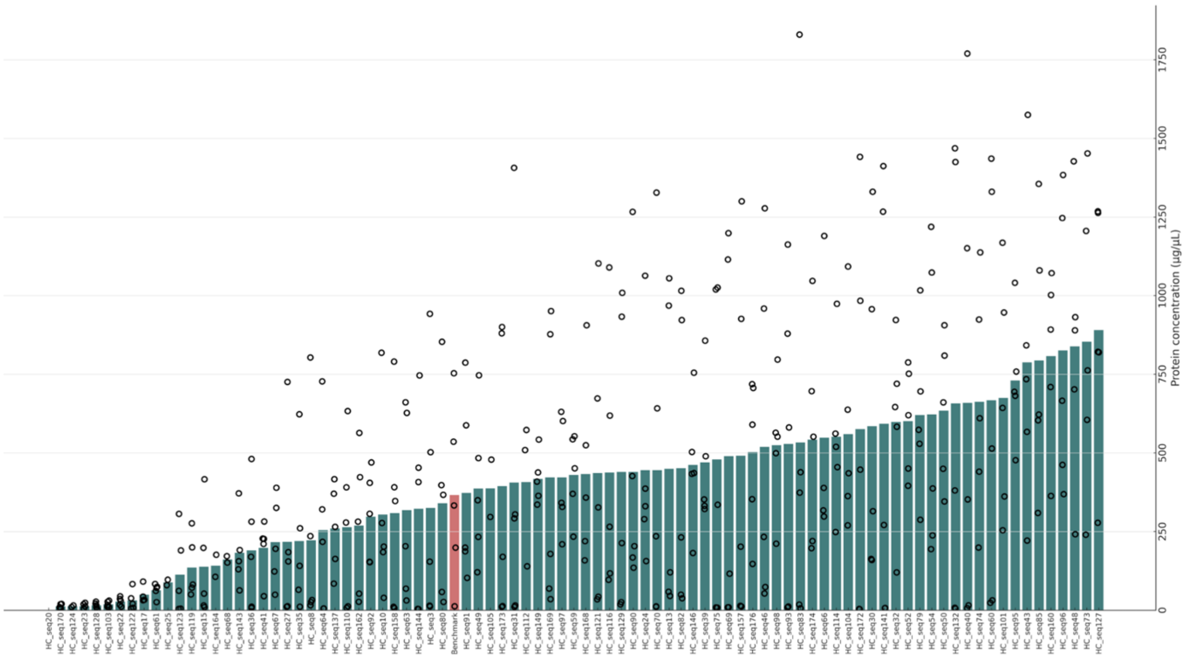


**Figure S5. Synthetic signal peptides lead to VHH-Fc secretion in the CHO G22 industrial cell line.** Protein concentration of the VHH-Fc in crude supernatant from CHO G22 cells after 10 days of expression in deep 96w plates measured through biolayer interferometry. The data is aggregated from 4 independent experiments comprising 5 technical repeats.
